# Optimization of Dose Schedules for Chemotherapy of Early Colon Cancer Determined by High Performance Computer Simulations

**DOI:** 10.1101/420232

**Authors:** Chase Cockrell, David E. Axelrod

## Abstract

Cancer chemotherapy dose schedules are conventionally applied intermittently, with dose duration of the order of hours, intervals between doses of days or weeks, and cycles repeated for weeks. The large number of possible combinations of values of duration, interval, and lethality has been an impediment to empirically determine the optimal set of treatment conditions. The purpose of this project was to determine the set of parameters for duration, interval, and lethality that would be most effective for treating early colon cancer. An agent-based computer model that simulated cell proliferation kinetics in normal human colon crypts was calibrated with measurements of human biopsy specimens. Mutant cells were simulated as proliferating and forming an adenoma, or dying if treated with cytotoxic chemotherapy. Using a high performance computer, a total of 28,800 different parameter sets of duration, interval, and lethality were simulated. The effect of each parameter set on the stability of colon crypts, the time to cure a crypt of mutant cells, and the accumulated dose was determined. Of the 28,800 parameter sets, 434 parameter sets were effective in curing the crypts of mutant cells before they could form an adenoma and allowed the crypt normal cell dynamics to recover to pretreatment levels. A group of 14 similar parameter sets produced a minimal time to cure mutant cells. A different group of 9 similar parameter sets produced the least accumulated dose. These parameter sets may be considered as candidate dose schedules to guide clinical trials for early colon cancer.

## Introduction

Colorectal cancer starts with the abnormal proliferation of mutant cells in colonic crypts. The population of mutant cells may continue to increase and form an adenoma, an early stage of colon cancer.^1^ The purpose of this project is to determine optimal chemotherapy dose schedules that eliminate these mutant cells before they can form an adenoma, and also the system retains crypt function.

There were an estimated 135,430 new cases of colon cancer in the United States in 2017, and an estimated 50,260 deaths.^2^ These deaths indicate that in spite of stool-based screening^3^, and colonoscopic screening for adenomas, both of which do reduce the risk of fatal colorectal cancer^4^, there is still a need for other methods of early detection. As new methods for detecting pre-malignant abnormalities come to fruition, possibly by blood tests^5^, the next need will be to determine the optimal therapies for such early lesions.

In order to determine optimal chemotherapy dose schedules for early colon cancer it is necessary to take into account the many possible values of three parameters, e.g. the duration of the dose, the interval between doses, and the lethal effect of the dose on mutant cells and on normal cells in the crypt. Values of these three parameters may vary over a wide range. For instance, the value of duration may be from hours to days, the interval between doses may be from days to weeks, and the lethality may kill 5% to 95% of mutant cells at each treatment cycle. Because of the large possible range of values of each of the three parameters, it has not been possible to instantiate a clinical trial that would compare all of the very many possible different conditions. However, computer simulations can be used to explore many dose schedules.

For the results of computer simulations to be reliable for finding optimal conditions, two criteria need to be satisfied. The model must be realistic, and the computer system must be able to explore a large number of different parameter sets.

We previously described a realistic computer model of cell dynamics in human colon crypts.^6,7^ The model utilizes the concept of agent-based modeling, where each biological cell is represented as an agent that has specified properties and interacts with other agents and with its environment. It has the advantage of producing emergent behaviors derived from each type of agent, such as stem cells, proliferating cells, differentiated cells, and rapidly dividing mutant cells. Then the emergent properties of the system of agents (biological cells in a crypt) in different environments (cytotoxic chemotherapy) could be generated in computer simulations. The model was calibrated with measurements of cell types in human biopsy specimens. The emergent behavior of the model reproduced several properties observed in biological crypts, including the ability of normal cell dynamics to recover from perturbations such as cytotoxic drug treatments that will eliminate mutant cells.

The second criterion for the results of computer simulations to be reliable is that the computer system needs to be able to explore a large number of different parameter sets. The original colon crypt model was written in the NetLogo application. The code for this model in this application was sufficient to simulate a few parameter sets, but not adequate to explore a large number of parameter sets necessary to determine which are optimum.^8^ In order to explore a large number of different chemotherapy dose schedules (duration, interval, lethality), the NetLogo code was ported to C++, and simulations were run on high performance computing platforms (Edison at NERSC and Beagle at the Computation Institute – University of Chicago). Such simulations were previously successful in exploring the large behavioral landscape of a different agent-based model.^9,10^

The goal of the current project was to explore the effect of many different dose schedules for their ability to eliminate mutant cells in an adenomatous colon crypt, and to determine a subset of dose schedules that are optimum. Each dose schedule had different set of parameter values for duration, interval, and lethality. The criterion of an optimum dose schedule is either one that eliminates mutant cells in the shortest time or one that has the least accumulated dose, and that retains normal crypt function.

We report on the results of 28,800 different simulated dose schedules. We found 14 dose schedules with the shortest time to eliminate mutants from an adenomatous crypt, and 9 dose schedules with the least accumulated dose.

## Methods

The agent-based colon crypt model, written in the application NetLogo, was previously described in detail.^6^ The model assumes that each cell’s probability to proliferate or to die is determined by its position in two gradients along the crypt axis, a divide gradient and a die gradient. A cell’s type, stem cell, proliferating cell, or differentiated cell is determined by its position in the divide gradient. A cell born near the bottom of the crypt moves up and is removed at the top where it undergoes apoptosis. Mutant cells are those that divide and die with a higher (though distinct) probability, than normal cells at the same spatial position in the colonic crypt.

The NetLogo model was calibrated using measurements of the number and variation of each cell type in histological sections of normal human colon biopsies. The behavior of the model was verified by its ability to reproduce the number and variation of each cell type, to undergo crypt fission by neutral drift, to have mutants proliferate to fill the crypt and form an adenoma, and to be robust and recover from perturbation from cytotoxic agents.^6,7^

In order to efficiently explore the chemotherapeutic parameter space, the the NetLogo crypt model^6^ (https://doi.org/doi:10.7282/T3KH0QKV) was ported to C++ (https://doi.org/doi:10.7282/T33X8B0H) using established methods.^9,11^ For cytotoxic chemotherapy, the duration of therapy was varied from 1 to 24 time-steps with increments of 1 step; the interval of therapy was varied from 2 to 48 steps with increments of 2 steps; and the lethality parameter (arbitrary units) was varied from a value of 1 to a value of 50 in increments of 1. In order to account for the effects of randomness in the model, fifty stochastic replicates were performed for each of the 28,800 cytotoxic chemotherapeutic dose schedules.

## Results

### Effective Dose Schedules

Colon cancer starts in a crypt with the abnormal proliferation of mutant cells to form an adenoma. Cytotoxic chemotherapy can kill dividing mutant cells, but may also kill dividing normal cells. Intermittent chemotherapy dose schedules were sought that would kill all mutant cells but allow normal cell dynamics to recover between doses and retain crypt function.

Normal cell types in the crypt include quiescent stem cells at the bottom of the crypt, proliferating cells near the bottom third, and differentiated cells in the top two-thirds. Cells move from the region of proliferating cells up to the region of differentiated cells at the top and are removed from the crypt. A dose of a cytotoxic drug will kill proliferating cells but spare quiescent stem cells. Quiescent stem cells are resistant to cytotoxic drugs because they have a very low probability of dividing. As the normal proliferating cells are killed, some quiescent stem cells become activated and produce additional proliferating cells. If the cytotoxic dose is of short duration, the number of proliferating cells, and their differentiated progeny, will increase to the pretreatment levels. Normal cell homeostasis with quasi-stationary cell dynamics will be restored, and the crypt will recover.

The goal of this project is to determine an optimal set of dose-schedules (parameter sets) (consisting of dose duration, interval between successive doses, and dose lethality) that will kill mutant cells in the crypt before they can proliferate and form an adenoma, and will allow restoration of the number of normal cells and recovery of crypt function.

In this context the terms Duration, Interval, and Lethality are defined as follows: Duration of the dose is the time during which cells in a crypt are exposed to the cytotoxic agent. The Interval between doses is the time from the beginning of one dose to the beginning of the next dose. Lethality is the factor that increases the probability that a cell will die above that determined by its position in the crypt. The lethality factor affects both normal cells and mutant cells. Mutant cells, in initial simulations, had a 1.16 X probability of dividing than normal cells at the same position in the crypt, and a 1.1 X probability of dying than normal cells at the same position in the crypt. Each specific treatment parameterization consisted of a dose schedule described by a parameter set of one value of Duration, one value of Interval, and one value of Lethality.

Dose schedules were simulated that included combinations of 24 different values of Duration, 24 different values of Interval, and 50 different values of Lethality, for a total of 28,800 parameter sets. Each set of parameters was simulated in 50 independent runs. The output of each parameter set included whether the crypt recovered and survived a time period of 1200 ticks, and if so, the time to cure the crypt of all mutants. Of the 28,800 different parameter sets, 434 (1.5 %) were effective, i.e. eliminated all mutant cells, and crypts recovered (https://doi.org/doi:10.7282/T37M0C92).

### Dose schedules with shortest time to cure

The average time to cure crypts of mutant cells was calculated over 50 stochastic replicates of each of the 434 effective dose schedules. There is a broad range of average cure times, from 2 time-steps to 263 time-steps (Figure 1A). One step equals approximately 4.5 human hours.^6^

**Figure 1.**
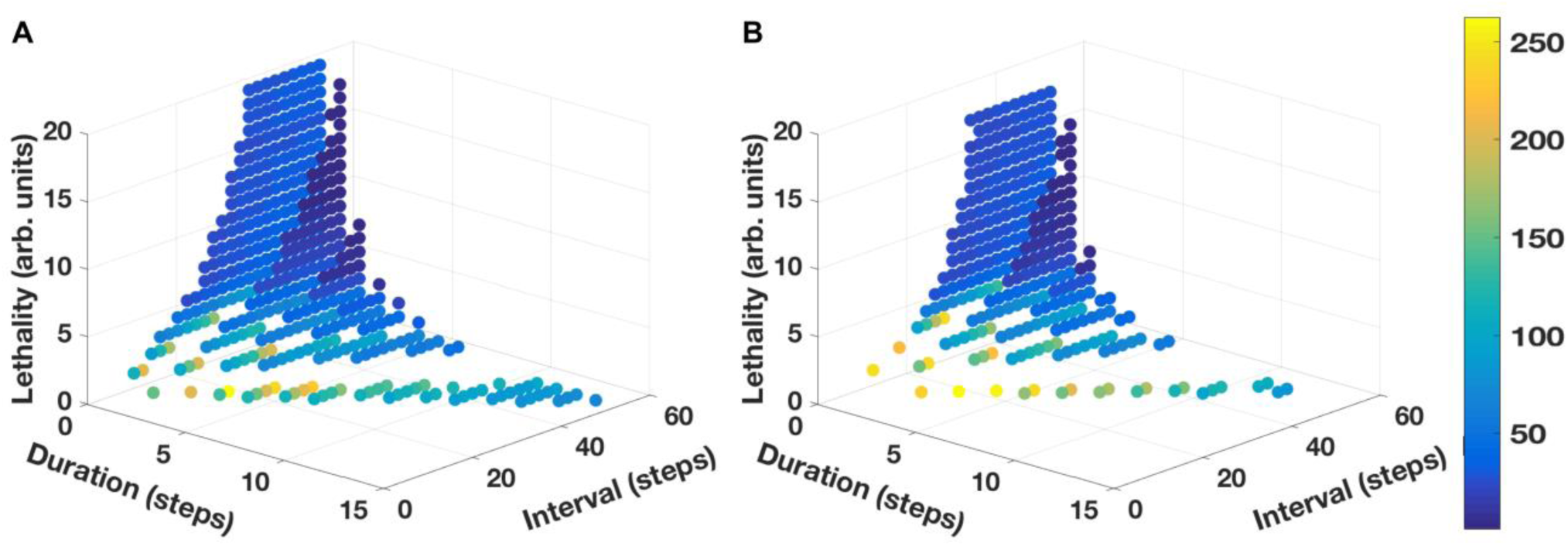
Distributions of average time to cure and sum dose of chemotherapy. Panel A: Time, in simulation steps, to cure mutants vs. the rank of each of 434 dose schedule parameter sets. Panel B: Sum of lethal doses to cure mutants vs. the rank of the each of the 434 dose schedule parameter sets. Points are color-coded according to the average time of 50 simulations that it takes to cure mutants, see Figure 3. Panel C: Comparison of the ranks of parameter sets for the sum of doses and the ranks of the parameter sets for the time to cure. The ranks are not are not highly correlated (R^2^ = 0.45).

Fourteen dose schedules had the shortest time to cure, 2 ticks. One of these is shown in Figure 2A. These dose schedules each had a Duration of 2 ticks, had Intervals of 42 to 48 ticks, and Lethality of 13 to 19. For comparison, the dose schedule that had the longest average cure time, 263 ticks, had a Duration = 4, Interval = 14, Lethality = 1 (Figure 2B). This indicates a range of interval/lethality combinations that have lethality sufficient to quickly kill all mutant cells, but an interval that is sufficiently long for the crypt to recover prior to the subsequent dose, thus establishing a dynamic equilibrium between the proliferative effects of the cells comprising the crypt with the cytotoxic effects of chemotherapy.

**Figure 2.**
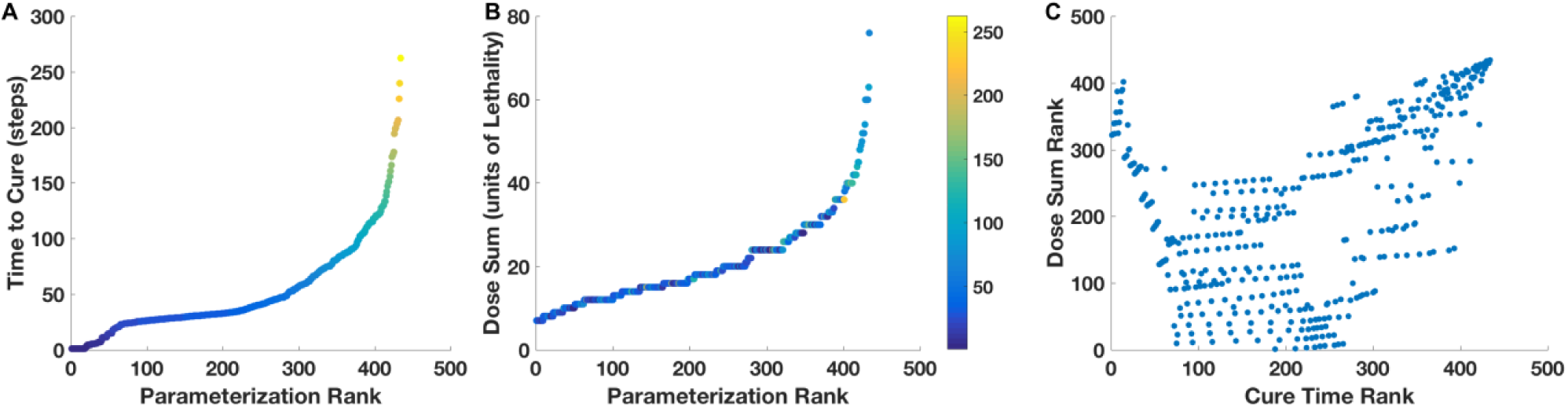
Time to cure and accumulated dose for different dose schedules. Panels A and B: Dose schedules for the shortest time to cure (2 time steps), and the slowest time to cure (263 time steps), with the indicated parameter set of Duration, Interval, and Lethality. The red arrows indicate the average times to cure for 50 simulations. Panels C and D: Example of one of 50 simulations of the kinetics of the proportion of mutants per crypt for the dose schedules shown in Panels A and B, respectively. Panels E and F: Total dose accumulation as a function of Time-steps for the dose schedules shown in Panels A and B, respectively. The average Total Dose at the time of cure is indicated by the red line.

The kinetics of the proportion of mutant cells per crypt as a function of time is shown for two example simulation runs, the shortest time to cure and the longest time to cure (Figure 2, C and D, respectively). Therapy was started after the proportion of mutant cells per total cells in the crypt was greater than 0.2, and the time to cure the crypt of all mutant cells was determined. The dose schedule with the shortest time to cure could eliminate all mutant cells within 2 ticks with only one dose, whereas the dose schedule with the longest time to cure required 20 successive intermittent doses to eliminate all mutants.

The decrease of the number of mutant cells as a function of time, and the recovery of the quasi-stationary number of each normal cell type in the crypt, can be seen in movies for the shortest time to cure (https://doi.org/doi:10.7282/T3RN3C79), and for the longest time to cure (https://doi.org/doi:10.7282/T3MW2MHK).

The 434 effective dose schedules can be visualized as a point in a three dimensional space with axes of Duration, Interval, and Lethality (Figure 3 A). The time to cure for each dose schedule is represented by a point of a different color. The group of dose schedules with shortest average cure time appear at a cusp on the surface of the three dimensional distribution. The arrangement of the different dose schedules in the three dimensional space can also be visualized as the figure is rotated in a movie (https://doi.org/doi:10.7282/T3CC141S 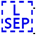).

**Figure 3:**
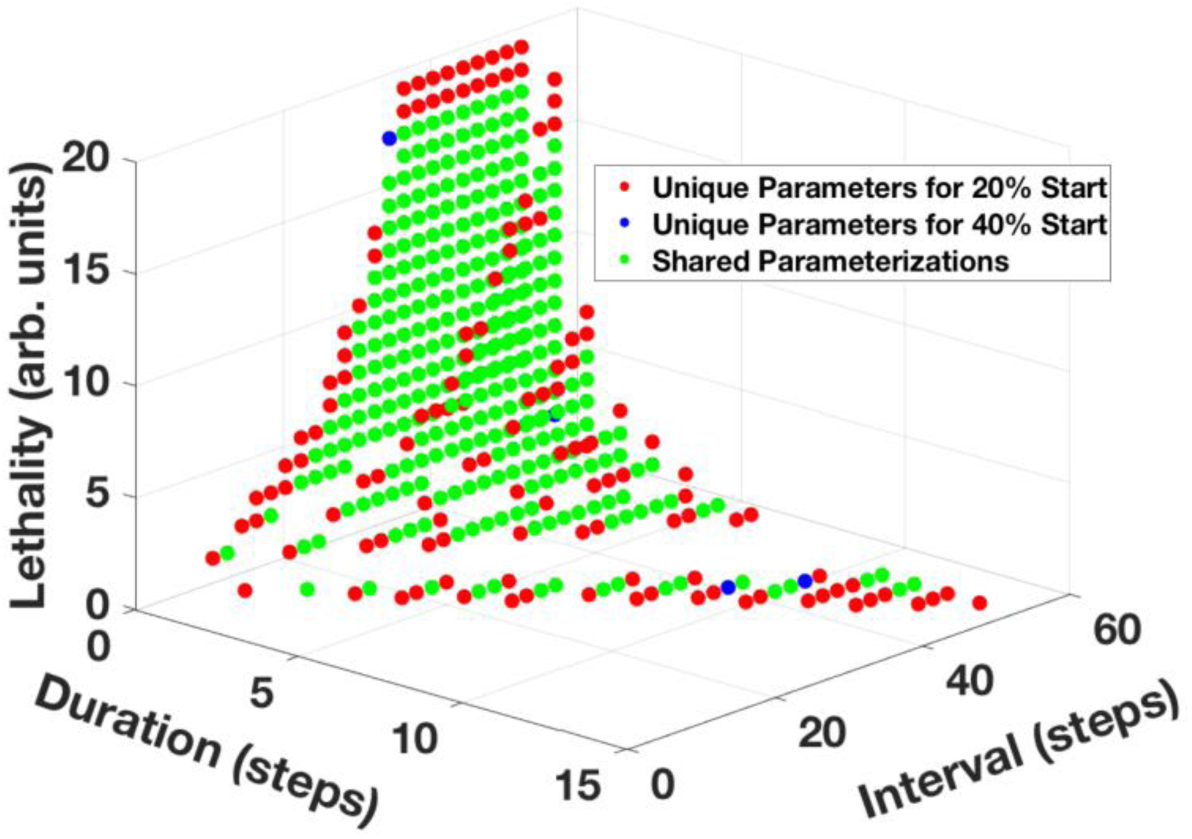
Dose schedule parameter sets that cure mutants when chemotherapy is started at different times. Each point represents a dose schedule parameter set that yield a 100% mutant cure rate and 0% crypt mortality rate when tested over 50 stochastic replicates. The Duration of chemotherapy (in simulation time-steps) is represented on the x-axis; the Interval between doses (in simulation time-steps) is represented on the y-axis; cytotoxic Lethality (arbitrary units) is represented on the z-axis. Points are color-coded based on the average time to cure a crypt of mutant cells, with red representing a treatment schedule that quickly eliminates the mutant cells and dark blue representing a treatment which takes longer to eliminate mutant cells. Panel A: Successful parameter sets for chemotherapy that is initiated when mutant cells make up 20% of the total crypt population; Panel B: Successful parameter sets for chemotherapy that is initiated when mutant cells make up 40% of the total crypt population. Panel C: Comparison of parameter sets in Panels A and B. Points shaded in red are those which are unique to the set of simulations that initiate chemotherapy when mutant cells compose 20% of the crypt; points shaded in blue are those which are unique to the set of simulations that initiate chemotherapy when mutant cells compose 40% of the crypt; points shaded in green are those which are shared between the set of simulations that initiate chemotherapy when mutant cells compose 20% of the crypt and the set of simulations that initiate chemotherapy when mutant cells compose 40% of the crypt.

### Mutants can be cured in simulations with different initial conditions

The effective cure times described above were determined for therapy that was started when the number of mutant cells was 20% of the total number of cells in the crypt. The mutants were a homogeneous population of cells that had a 1.16X probability of dividing and a 1.1X probability of dying compared to non-mutant cells at the same position in the crypt. These values are not unique; cures can be achieved with other values. As examples, we give results with three different sets of values.

First, the treatments were started when the proportion of mutant cells per crypt was twice, i.e. 40% rather than the 20% described above. Cures were also obtained when treatments were started at 40%. The distribution of cure times for doses started at 20% and 40% are shown in Figure 3, A and B, respectively. The similarities and differences in the cure times are shown in Figure 3 C.

Second, mutant cells were considered that grew half as fast, having half the probability of dividing, i.e. 1.08X rather than 1.16X of that of normal cells. Cures were also obtained with the slower growing mutant cells. Slower-growing mutants would be expected to be more resistant to cytotoxic chemotherapy than the faster growing mutants. The average cure time of the slower growing mutants was 17 ticks rather than 2 ticks of the faster growing mutants.

Third, a heterogeneous population of mutant cells, rather than a homogeneous population, was considered. Biological crypts may be heterogeneous, containing different cells with different mutations and different drug responses.^12^ Cures were also obtained with a heterogeneous population of mutant cells. A heterogeneous combination of fast and slow growing mutants had an average cure time of 12 ticks, rather than a cure time of 2 ticks for the homogeneous population of faster growing mutants.

Therefore, mutant cells can be cured under different initial conditions, e.g. when treatment is started when there are different proportions of mutant cells, when the mutant cells are relatively fast or slow growing, and when the population of mutant cells is homogeneous or heterogeneous.

### Dose schedules with least treatment to cure

Repeated cycles of treatments (Figure 2, A and B) result in the accumulation of doses over time (Figure 2,E and F). Where each Dose = Lethality x Duration, and the accumulated dose is the sum of Doses up to the time that all mutants are cured. The least amount of accumulated dose needed to eliminate mutant cells may be considered as an alternative criterion for optimal therapy to the shortest time to eliminate mutant cells.

The distribution of accumulated doses to cure for each of the 434 effective treatment schedules is shown in Figure 1,B. This distribution appears to be similar to the distribution of least time to cure (Figure 1,A). However, the ranks of the parameter sets for accumulate doses to cure and ranks of the parameter sets for the least times to cure are not highly correlated (R^2^ = 0.45) (Figure 1,C).

It should be noted that the parameter set of the dose schedule with the least accumulated dose that cures mutants, is not the same as the parameter set of the dose schedule with the sum of doses with the shortest time to cure (Figure 1, C and Table 1). Therefore, the least accumulated dose to cure and the shortest time to cure, are two independent criteria for optimal chemotherapy.

**Table 1.** Comparison of Schedules with Shortest Cure Time and Least Treatment

| Criterion | Avg Cure Time | Dose Sum | Duration | Interval | Lethality |
| --- | --- | --- | --- | --- | --- |
| Shortest time | 2 | 26 | 2 | 42 | 13 |
| Least treatment | 32 | 7 | 1 | 32 | 7 |

Note: Where 1 tick = 4.5 hr, and Dose = Lethality x Duration.

## Discussion

In this project, many different intermittent dose schedules of a cytotoxic chemotherapeutic drug have been evaluated for their ability to eliminate mutant cells from a colon crypt before the mutants can fill the crypt and form adenoma. Eliminating mutant cells, while retaining crypt function, could intercept the progression of adenomas to adenocarcinomas.^13,14^

Each different dose schedule consisted of a different set of values of duration of the dose, interval between doses, and lethality of the dose. A high performance computer was used to simulate the effect of 28,800 different sets of dose schedules. Of these, a subset of 434 dose schedules was effective, e.g. they eliminated all mutant cells, and allowed crypt normal cell dynamics to recover from the lethal effects of the chemotherapy. The other dose schedules were not effective; although they eliminated mutant cells, they also killed most normal cells, resulting in total collapse of the crypt. Effective dose schedules were identified that produced the shortest time to cure all mutants from the crypt, and different effective dose schedules were identified that produced the least accumulated dose.

The ability of a subset of intermittent dose schedules to both eliminate mutant cells and retain normal cell crypt function was shown to be effective for a range of initial conditions, including the following: (1) drug treatment could be initiated when the mutants are 20% or 40% of the total number of cells in the crypt, (2) mutants could be eliminated that proliferated much faster than normal cells and are relatively sensitive to cytotoxic chemotherapy, and mutants could be eliminated that proliferate only slightly faster than normal cells and are relatively resistant to chemotherapy, and (3) the mutant populations may be homogeneous, or may be heterogeneous consisting of both fast and slow proliferating mutant cells.

Cell dynamics in the human normal and neoplastic crypts has been the subject of other modeling studies.^15,16^ This is partially due to the fact that cell dynamics have been well characterized experimentally.^17^ In a normal unperturbed crypt, homeostasis is maintained with a quasi-stationary number of total cells, and of each cell type. The average total number of cells has been measured in human colon crypts is 2,427. The average number of quiescent stem cells, proliferating cells, and differentiated cells is 36, 624, and 1768, respectively.^6^

Stem cells in the normal crypt, may be quiescent or may become active^18^, and divide stochastically.^19,20^ Actively dividing stem cells yield proliferating cells, which in turn move up the crypt, differentiate, and are removed at the top.^21^ The rate of cell loss at the top of the crypt is balanced by the rate of stem cell division at the bottom of the crypt. This balance maintains homeostasis.^22^ The rate of stem cell divisions is controlled by the number of stem cells, by the number of proliferating cells, and indirectly by the number of differentiated cells.^23^

If a crypt is exposed to a brief low dose of a cytotoxic chemotherapeutic drug, some proliferating cells are killed and the stem cells will respond by producing more proliferating cells. This results in restoration of homeostatic cell dynamics and recovery of crypt function. If mutant cells are also in the crypt, a longer or a more lethal dose may be required to kill the mutant cells before they can proliferate and fill the crypt to form an adenoma. However, such a longer or more lethal cytotoxic dose may overwhelm the ability of the stem cells to repopulate the crypt and the crypt will collapse.

Cell dynamics in the colon crypt without therapy have been modeled mathematically^24^ and computationally^6,25^, and reviewed.^26,27,28^ Crypt dynamics with therapy has been modeled in order to determine a therapy that could eliminate mutant cancer cells but still allow recovery of normal crypt cell homeostasis. Optimal control theory for cancer therapy has been reviewed^29,30^, and computer models for cancer therapy have been reviewed.^31^

Many of the published modeling reports provide a different perspective than the modeling results that we report. We focused our attention on an early stage of colon cancer initiated by abnormally proliferating mutant cells in a colon crypt that also had normal cells. Our agent-based computer model was calibrated with measurements of human biopsy specimens. We simulated the response to cytotoxic chemotherapy of both mutant cells and normal cells. By comparison, Panetta^32^ described a competition model with periodically pulsed chemotherapy and parameter values needed to prevent relapse. Gaffney^33^ mathematically modeled schedules with rest phases between chemotherapy and emphasized the importance of choosing the correct intervals between doses. Marcu and Bezak^34^ used Monte Carlo computer modeling in order to determine the conditions under which intermittent therapy may fail because some tumor cells repopulation the tumor between doses. Murano et al. ^36^ mathematically modeled normal crypt cell dynamics and concluded that stem cells and proliferating cells should react differently to therapy-induced apoptotic killing. Leder et al.^36^ mathematically modeled radiation dose schedules and predicted that hyper-fractionated dosing schedules would be superior to hypo-fractionated dosing schedules. These results were confirmed for survival of irradiated mice with glioblastoma.

We have considered two kinds of optimal dose schedules, in each case allowing recovery of normal cell dynamics and crypt function. One kind of optimal schedule eliminates all mutant cells in the shortest time, and a second kind of optimal schedule eliminates all mutants with the least accumulated dose. A choice may be made between these two optimal schedules in order to reduce the collateral damage due to chemotherapy, such as neuropathy, cardio-toxicity, and neuropenia. Alternatively, rather than choose one or the other, there are several parameter sets that have both intermediate times to cure and intermediate accumulated doses (Figure 2 C). These parameter sets provide an opportunity to select a dose schedule treatment that is good, but not perfect, by each criterion.

We acknowledge several limitations of this project. For instance, we have not taken into account that cytotoxic drugs may induce new mutant cells, including drug resistant mutants, as well as kill existing mutant cells. However, we have already modeled the induction of new mutant cells and shown that intermittent dose schedules can be effective in eliminating mutants that arise spontaneously or are induced by a cytotoxic drug^7^. Also, this project has not modeled chemotherapy dose schedules by a combination of two, or more, drugs. Such an extension will require considering duration, interval, and lethality of at least two drugs, and different orders of the different drugs. Simulating, and evaluating the output of such a large number of sets of parameters, and large range of values of each parameter, will require machine learning strategies for multi-object parameterization^9,37^ rather then the parameter sweeping strategy that was sufficient for simulating a single drug.

In conclusion, 28,800 sets of intermittent chemotherapeutic dose schedules for early colon cancer were evaluated for their ability to remove all mutant cells while retaining normal crypt function. Each dose schedule had a different duration of chemotherapy dose, different intervals between doses, and different lethality. A subset of similar dose schedules was determined that had the shortest time to cure and another subset of similar dose schedules was determined that had with the least accumulated dose. These subsets of effective dose schedules suggest candidate dose schedules for clinical trials.

## Acknowledgements

C.C. thanks Gary An for discussions on agent-based modeling techniques. D.E.A. thanks the members of the Division of Life Sciences IT Support Group for computer services, the RUCore staff for file archive services, and Albertas Janulevicius for suggestions on how to increase NetLogo executions speed, which lead to the collaboration with C.C.

## Author’s contributions

C.C. ported D.E.A.’s model NetLogo code to C++, and ran the simulations. D.E.A. wrote a first draft. Both authors produced graphs, interpreted data, and approved the final draft.

